# Directing iPSC Differentiation into iTenocytes using Combined Scleraxis Overexpression and Cyclic Loading

**DOI:** 10.1101/2021.11.23.469329

**Authors:** Angela Papalamprou, Victoria Yu, Angel Chen, Tina Stefanovic, Giselle Kaneda, Khosrowdad Salehi, Chloe Castaneda, Arkadiusz Gertych, Juliane D Glaeser, Dmitriy Sheyn

**Author notes:** Correspondence: Dmitriy Sheyn, PhD Assistant Professor, Board of Governors Regenerative Medicine Institute Department of Orthopedics, Department of Surgery, Department of Biomedical Sciences Cedars-Sinai Medical Center, 8700 Beverly Blvd. AHSP A8308 Los Angeles, CA, 90048, USA.

## Abstract

Regenerative therapies for tendon are falling behind other tissues due to the lack of an appropriate and potent cell therapeutic candidate. This study aimed to induce cell tenogenesis using stable Scleraxis (Scx) overexpression in combination with uniaxial mechanical stretch of mesenchymal stromal cells (MSCs) of different origins. Scleraxis (Scx) is the single direct molecular regulator of tendon differentiation known to date. Mechanoregulation is known to be a central element guiding tendon development and healing. Cells explored were bone marrow-derived (BM-)MSCs as well as MSCs differentiated from induced pluripotent stem cells (iMSCs). Mechanical stimulation combined with Scx overexpression resulted in morphometric and cytoskeleton-related changes, upregulation of early and late tendon markers, increased ECM deposition and alignment, and tenomodulin perinuclear localization in iMSCs, which was greater compared to BM-MSCs and controls. Our findings suggest that these cells can be differentiated into tenocytes and may be a better candidate for tendon cell therapy applications than BM-MSCs.

## 1. Introduction

Tendon and ligament injuries are the main reason for all musculoskeletal consultations worldwide and represent a significant concern in sports medicine, as well as the general population. An estimated 30 million tendon and ligament cases are reported annually worldwide.^1^ In the US alone, more than 100,000 tendon and ligament repair surgeries are performed annually, resulting in more than $30 billion expenditure.^2^ Tendon injuries are associated with a high morbidity, prolonged disabilities, and painful rehabilitation periods, while secondary tendon ruptures often ensue. Current treatment modalities include autografts, allografts, xenografts, suture techniques, and prostheses.^2;3^ These options have major disadvantages including donor site morbidity, risk of limited long-term function, and an incidence of osteoarthritis in up to 50% of young patients.^2; 4^ Following treatment, functional recovery of repaired tendons is usually incomplete. The mechanical and structural properties of repaired tissue are permanently altered and fail to reach the level of functionality achieved prior to injury.^5^ Development of a cell therapy approach, which could be applied to an injured tendon or ligament site, could dramatically alter patient outcomes.

Adult stem/progenitor cells that have been identified in the tendon niche and characterized by their marker profile are considered promising cell sources for tendon tissue engineering.^6^ However, they are very scarce *in vivo* and cannot be expanded *in vitro* for therapeutic applications due to phenotypic drift in culture.^7^ Thus, adult stem/progenitor cell-mediated therapies have shown limited potential to be used for tendon repair.^8–10^ In addition, mesenchymal stromal cells (MSCs) derived from bone marrow (BM-MSCs) or adipose tissue (ASCs) have been explored as a potential tendon repair and tissue engineering strategy due to their abundance, multipotency, and regenerative potential *in vivo.* Several animal models have shown improved functional outcomes and enhanced tendon healing via utilizing MSCs.^8; 9; 11^ However, their major disadvantages are their limited self-renewal capacity, phenotypic heterogeneity, and potential for ectopic bone or cartilage formation. These attributes may limit their clinical application due to the need for *in vitro* expansion in adequate numbers and thorough characterization to meet regulatory standards prior to clinical use.^5^ Additionally, their tenogenic potential has been shown to be restricted.^12^ Developing an off-the-shelf cell source that can be efficiently differentiated to tenogenic progenitor cells is a prerequisite for tendon cell-based therapy success.

The discovery of induced pluripotent stem cells (iPSCs) through nuclear reprogramming of somatic cells revolutionized the field of regenerative medicine, because of their high self-renewal capacity and unparalleled developmental plasticity. Additionally, iPSCs might have the advantage of being able to evade the recipient’s immune response.^13; 14^ iPSCs have been successfully differentiated to MSC-like cells (iMSCs) by our group and others.^15; 16^ One of the main advantages of using iMSCs is that they potentially represent an unlimited source for tenocytes. Rodent tendon repair models utilizing iPSCs have shown improved functional outcomes.^17; 18^ Thus, iMSCs derived from iPSCs may offer an unlimited off-the-shelf allogeneic source for tendon cell therapy applications.

Development of tenocytes occurs in at least two stages: First, tenocyte progenitors (tenoblasts) express scleraxis (Scx), which is the single direct molecular regulator of tendon differentiation known to date.^19; 20^ Second, tenocyte maturation results in tissue formation.^20^ However, ectopic overexpression of Scx in BM-MSCs was not found sufficient to drive tenogenesis.^19; 21^ Tendon and other tissues grow and remodel in response to changes in their environment. At the cell level, spatial distribution of dynamic mechanical cues can affect developmental, maintenance, and healing responses.^22; 23^ Mechanoregulation has been shown to be a central element guiding embryonic tendon development and healing.^24^ Fully differentiated cells, such as tenocytes and fibroblasts, can actively sense both the external loading applied to them and the stiffness of their environment.^24; 25^ Mechanical cues can also affect the differentiation of multipotent cells such as BM-MSCs.^26; 27^ Additionally, tenocytes and tendon progenitors are able to retain their phenotype when they are stretched or cultured on mechanovariant substrates with controlled mechanical properties.^28^ Uniaxial cyclic stretching has been used to drive tenogenic differentiation *in vitro,* as it is considered more relevant physiologically.^29^ Therefore, mechanical stimulation may be essential for the differentiation of multipotent cells towards tenogenic progenitors, as well as for the maturation of the tenocyte phenotype.

Tenocytes can actively sense external loading, retain their phenotype when stretched, and dedifferentiate when cultured statically. Additionally, Scx is a crucial factor for tenogenic differentiation.^7; 19; 20^ Therefore, combinatory mechanical and biological stimulation may be essential for tenogenesis. We hypothesized that iMSCs can be efficiently differentiated into tenocytes by combination of stable overexpression of Scx and mechanical stimulation *in vitro*. In this study we investigated (a) the ability of stable Scx overexpression to induce tenogenic differentiation, and (b) the effect of mechanical stimulation to guide tenogenic differentiation *in vitro* with and without Scx overexpression. To accomplish this, Scx was overexpressed using lentiviral vectors. BM-MSCs, BM-MSC^SCX+^, iMSCs, and iMSC^SCX+^ cells were grown on mechanovariant substrates. Cyclic uniaxial stretching was applied on iMSCs and iMSC^SCX+^ using the CellScale MCFX bioreactor system. The effect of Scx overexpression with and without mechanical stimulation was assessed with gene expression analyses and immunocytochemistry for tendon markers, collagen deposition, and morphometric analysis of cytoskeletal orientation following uniaxial stretching.

## 2. Methods

### Primary cell isolation and expansion

Human bone marrow was acquired from Lonza (Allendale, NJ) and BM-MSCs were isolated as previously reported.^30–32^ Briefly, bone marrow was overlayed on Ficoll-paque density gradient medium (GE healthcare, Chicago, IL), washed twice with phosphate buffered saline (PBS), plated at a density of 2×10^5^/cm^2^ (Corning, Corning, NY), and incubated overnight at 37°C/5% CO_2_. After 48h, non-adherent cells were discarded, washed with phosphate buffered saline (PBS), and culture medium was added, composed of 2mM L-glutamine, 1% antibiotic antimycotic solution (AAS) in Dulbecco’s modified eagle medium (DMEM, all from Thermofisher, Waltham, MA), and 20% fetal bovine serum (FBS, GeminiBio, West Sacramento, CA). BM-MSCs were isolated via adherence and cultured at 37°C/5% CO_2._ Media were switched to standard MSC culture media (low glucose DMEM, 1% AAS, 10% FBS, 2mM L-glutamine) and were changed every 3 days. Cells were split upon confluence in a 1:3 ratio.

Primary porcine tenocytes were isolated from pig Achilles tendons. Briefly, tendons were washed with PBS containing antibiotics, manually minced to 2-3mm fragments, and digested with 0.02% collagenase II (LS004205, Worthington Biochemical Corporation, Lakewood, NJ) for 18h at 37°C/5% CO_2_ with rocking. The next day, pre-warmed media were added (high glucose DMEM supplemented with 1% AAS, 10% FBS, and 2mM L-glutamine), and tissue digests were centrifuged and strained through a 70 μm filter. Isolated cells were cultured in high glucose DMEM supplemented with 1% AAS, 10% FBS, and 2mM L-glutamine.

### iMSC derivation and expansion

Normal human iPSC lines were obtained from the Cedars-Sinai iPSC core facility, expanded on Matrigel^TM^-coated plates (Corning) with chemically defined mTeSR™Plus media (StemCell Technologies, Vancouver, Canada). iMSCs were differentiated from human iPSCs and expanded *in vitro* using our previously published method.^15^ iMSCs were maintained following the same protocol and media as described for BM-MSCs. Media were changed every 3 days, and cells were split upon 80% confluence in 1:3 ratio.

### Genetic engineering of BM-MSCs and iMSCs to overexpress Scx

BM-MSCs and iMSCs were engineered via lentiviral transduction to overexpress Scx-GFP under the constitutively active CMV promoter coupled to its enhancer using a lentiviral vector (BM-MSC^Scx-GFP+^, iMSC^Scx-GFP+^).^33; 34^ Briefly, HEK293T/17 cells (ATCC) were seeded at a density of 6×10^4^/cm^2^ in Eagle’s Minimum Essential Medium (EMEM, Thermofisher, Waltham, MA), containing 10% FBS, and 1% AAS. Lentiviruses were produced by transfecting HEK293T/17 cells with the pLenti-C-mGFP vector where SCXB has been inserted (OriGene, Rockville, MD) expressing Scx-GFP and two packaging plasmids^33^ (pCMV-dR8.2, pCMV-VSV-G). Transfections were carried out using the BioT method with a 1.5:1 ratio of BioT (µl) to DNA (µg). The virus-containing medium was harvested at 48 and 72h after transfection. Both batches were combined, centrifuged, filtered, and used to transduce BM-MSCs and iMSCs within 24h. Transduction titers were determined using flow cytometry for GFP and verified using RT-qPCR for Scx.

### Flow cytometry

For assessment of lentiviral transduction efficiency, transduced cells were washed with PBS and digested with 0.05% Trypsin-EDTA (Gibco) for 5 min at 37°C. Cells were then washed with FACS buffer containing 2% bovine serum albumin (BSA, A4503, Sigma, St. Louis, MO) and 0.1% sodium azide (S2002, Sigma) in 1x PBS, and were acquired on a BD LSR Fortessa analyzer (BD Bioscences, San Jose, CA). Generated FSC files were analyzed using Flowjo software (Flowjo LLC, Ashland, OR).

### Cell culture on various stiffness surfaces

BM-MSCs and iMSCs were seeded at 2.5×10^4^/cm^2^ in plates of varying stiffness. Specifically, cells were seeded into 6-well Cytosoft^®^ (Advanced Biomatrix, Carlsbad, CA) plates which are pre-coated with polydimethylsiloxane (PDMS) by the manufacturer with (E) of E~2kPa and E~32kPa, as well standard tissue culture polystyrene (TCP) plates (E~GPa range) (Corning). Prior to seeding the cells, PDMS plates were coated with Collagen-1 (Purecol^®^, Advanced Biomatrix), following the manufacturer’s recommendations. Cells were grown 12 days in tenogenic media (low glucose DMEM supplemented with 2mM glutamine, 10% FBS, 1% AAS, and 50µg/ml ascorbic acid).^35; 36^ After 12 days, cells were lysed with RLT plus lysis buffer (RLT plus buffer, Qiagen, Valencia, CA containing β-mercaptoethanol, M3148, Sigma, St. Louis, MO), and stored at −80°C until processing for gene expression analysis.

### Mechanical loading

Since tendon cells predominantly receive uniaxial tensile loading from collagen bundles *in vivo*,^29^ the CellScale MCFX bioreactor (CellScale Biomaterials Testing, Waterloo, ON, Canada) was chosen, as it allows for longitudinal stretching on the cell monolayer. Cells were seeded at a density of 2×10^4^/cm^2^ on fibronectin-coated silicone plates (CellScale Biomaterials Testing) and allowed to attach for 24h. Since the strain gradients in developing tissues are not well-known, the cyclic stretching protocol was optimized based on published literature at 4% sinusoidal strain, 0.5 Hz, 2 h/day.^37; 38^ Cells seeded on the same fibronectin-coated silicone plates were placed in the incubator in parallel and served as static controls for those experiments. iMSCs and iMSC^Scx+^ were stretched for up to 7 days, and media were changed 3x/week in tenogenic media (also used for *in vitro* static culture). Cells were harvested on days 0, 3, and 7 (n=8/timepoint).

### Cell morphology and morphometric analysis

At all time-points (days 3 and 7), cells were fixed with 10% buffered formalin for 15 min at RT, permeabilized in PBS with 0.3% Triton X-100 (PBS-T) and stained with anti-F-actin iFluor 555 phalloidin antibody (Abcam, Cambridge, UK) to identify the cytoskeleton. Nuclei were counterstained with DAPI (Thermofisher). Samples were imaged with an Inverted Revolve fluorescent microscope (Revolve, Echo, San Diego, CA) with a 200x magnification objective using a 3×3 grid to capture 9 non-overlapping images for analysis. ImageJ was used to quantify Tenomodulin (Tnmd) staining intensity in unprocessed single channels to extract relative pixel intensity per field of view. No primary antibody-treated samples (secondary antibody included) were used to normalize this analysis. Lastly, to quantify organization of the cytoskeleton in each cell, the ImageJ directionality tool was used to assess actin filament angle distributions.

Single-cell morphometric analysis was carried out using a custom image analysis pipeline developed in MATLAB (Matlab Natic, MA), as previously described.^39^ The analysis was performed in unprocessed single channels by a bio-image analysis expert, who was blinded to the image groups. First, nuclear mask was obtained from the DAPI image ^40^. Based on this mask, the area, orientation cell density, and aspect ratio (width:height of the nucleus) were measured.^39^ Next, the number of nuclei in the mask was divided by the image area to determine cell density per field of view. For this analysis n=7 wells/group with 5 non-overlapping high-powered fields of view per well were used. Nuclear orientation and ImageJ directionality data were displayed as relative frequency distribution rose plots using MATLAB, with a bin width of 5° and 10°, respectively. A nuclear orientation or actin direction of 90° indicates that the nucleus or actin fiber are angled perpendicular to the direction of stretch.

### Gene expression analysis

Differentiation to iPSC-derived Tenocytes (iTenocytes) was defined based on expression of tendon-specific markers (Suppl. Table 1) using RT-qPCR with TaqMan^®^ gene expression as well as osteogenic and adhesion markers.^41; 42^ Total RNA was isolated to analyze the gene expression of tendon phenotypic markers (Suppl. Table 1). For RNA isolation, media were aspirated and cells were lysed with RLT plus lysis buffer (Qiagen). Lysates were transferred to 1.5 ml tubes and were fully homogenized using a handheld pestle and mortar. RNA was isolated using the RNeasy plus mini kit (Qiagen) and RNA yields were determined spectrophotometrically using a Nanodrop system (Thermofisher). RNA was reverse-transcribed with the high-capacity cDNA reverse transcription kit (Applied Biosystems). Target gene mRNA levels were quantified using FAM-MBG technology (Bio-Rad, Hercules, CA). The threshold cycle (Ct) value of 18S rRNA was used as an internal control using the TaqMan^®^ gene expression FAM/MGB probe system (4333760F, Thermofisher). The Livak method was used to calculate ΔΔCt values and fold change was calculated as 2^−ΔΔCt^, as previously described and published.^30; 31; 43–46^

### Assessment of collagen deposition

Newly synthesized collagen was quantified using the Sirius red total collagen assay detection kit (Chondrex, Redmond, WA). Total collagen in each cell culture media sample was concentrated using Concentrating Solution (Chondrex) at 4°C for 18-24h. Following incubation, samples were centrifuged for 3min at 10,000 rpm. Briefly, collagen pellets were dissolved in 0.05M acetic acid (Sigma), precipitated with Sirius Red dye, washed, and then transferred to a 96-well plate. Optical density at 535 nm was determined for samples and assay standards. Collagen concentration was calculated from the standard curve using regression analysis and plotted in µg/ml units. All experimental samples (n=8) and assay standards were conducted in technical triplicates.

### Immunocytochemistry (ICC)

Cell cultures of iMSC^SCX+^ were grown in static culture or uniaxially stretched in the 2D bioreactor for 7 days as described above. After 7 days, cells were briefly washed with PBS and fixed with 10% buffered formalin for 15 min at RT. Cells were washed again, and nonspecific sites were blocked with 10% normal donkey serum (Jackson Immunoresearch, West Grove, PA) in 0.3% v/v PBS-Tween® 20 Detergent (PBS-T) for 45 min. After, cells were incubated at 4°C overnight with the primary antibody for Tnmd (HPA055634, Sigma, Darmstadt, Germany) or Collagen-1 (1310001, Bio-Rad). Cells were then washed 3 times with PBS-T, followed by a 2h incubation at RT with the secondary antibody donkey anti-rabbit Alexa Fluor^®^ 647-conjugated Affinipure (Jackson Immunoresearch). Nuclei were counterstained with DAPI. Fluorescent images were captured with a Revolve fluorescent microscope (Revolve). Images of single channels were taken, and they were merged either during imaging or using Adobe Photoshop CS6 post imaging. Adobe Illustrator CS6 was used for final figure production. BioRender software program was used to prepare the graphical abstract and additional graphics used throughout the manuscript.

### Statistical Analysis

All data are presented as mean ± standard deviation from the mean. Normally distributed data were analyzed with unpaired t-test (for 2 groups), or non-repeated measures analysis of variance followed by Tukey-Kramer HSD *post hoc* analysis when more than 2 groups were compared. Non-parametric data were analyzed using the Mann-Whitney and Kruskal-Wallis tests. To compare gene expression levels over time, a mixed repeated measures model was used, followed by Tukey’s post hoc tests for between group comparisons. Lastly, to analyze the orientation of nuclei and actin filaments between the static and stretched conditions, medians were compared using the Mann-Whitney test, while the two frequency distributions were compared using the Kolmogorov-Smirnov test. Statistical significance was set at p<0.05. Statistical analyses were performed with GraphPad Prism 9.0.

## 3. Results

### 3.1 Confirmation of stable lentiviral-driven Scx overexpression in BM-MSC^SCX+^ and iMSC^SCX+^

We generated lentiviral vectors expressing Scx tagged with GFP at the C-terminus and under the constitutively active CMV promoter, as previously described.^33; 34^ The absolute intensity of GFP-positive cells was used as a proxy for Scx integrations using flow cytometry. Absolute GFP fluorescence was similar for the highest titers (Suppl. Fig. 1A). A dose-response effect of lentiviral load in transduction efficiency was observed when flow results were presented as percentage of GFP+ cells (Supp Fig. 1B). We expanded BM-MSCs and iMSCs transduced with Scx-GFP lentivirus vector (BM-MSC^SCX+^ and iMSC^SCX+^) and assessed the expression of Scx at 4 weeks of regular culture without sorting. Scx expression was significantly upregulated in BM-MSC^SCX+^ and iMSC^SCX+^ showing stable overexpression of Scx after 4 weeks (Suppl. Fig. 1C).

### 3.2. Tenogenic markers gene expression of BM-MSCs, BM-MSC^SCX+^, iMSCs, and iMSC^SCX+^ after 12 days of static culture

First, we examined the effect of Scx overexpression on BM-MSCs and iMSCs *in vitro*. The four cell groups BM-MSCs, BM-MSC^SCX+^, iMSCs, and iMSC^SCX+^ were generated and cultured *in vitro* on standard TCP culture plates for a duration of 12 days (Fig. 1A). Findings were summarized in a heat map showing changes compared to baseline (Fig.1B) and as bar graphs displaying differences between all cell types (Fig. 1C). Tenogenic marker genes (Scx, Mkx, Thbs4, and Tnmd) and ECM proteins previously associated with tenogenesis (Bgn, Col3a1, and Dcn) were significantly upregulated in iMSC^SCX+^ compared to their own baseline (day 0) whereas in BM-MSC^SCX+^ only Scx, Bgn, Thbs4, and Dcn were significantly upregulated compared to their own baseline (Fig. 1B). Mkx and Tnc showed significant downregulation in BM-MSC^SCX+^ and in BM-MSCs. Scx was demonstrated to be significantly overexpressed in BM-MSC^SCX+^ and iMSC^SCX+^ compared to BM-MSCs and iMSCs after 12 days (Fig. 1C). When compared to other cell types, in the iMSC^SCX+^ group, expression of Mkx, Bgn, Thbs4, and Tnmd was significantly higher. Thbs4 was significantly higher in BM-MSC^SCX+^ versus BM-MSCs. However, Tnmd was significantly higher in BM-MSCs compared to BM-MSC^SCX+^. Dcn was significantly upregulated in iMSC^SCX+^ but was lower compared to both BM-MSCs and BM-MSC^SCX+^ (Fig. 1C).

**Fig. 1.**
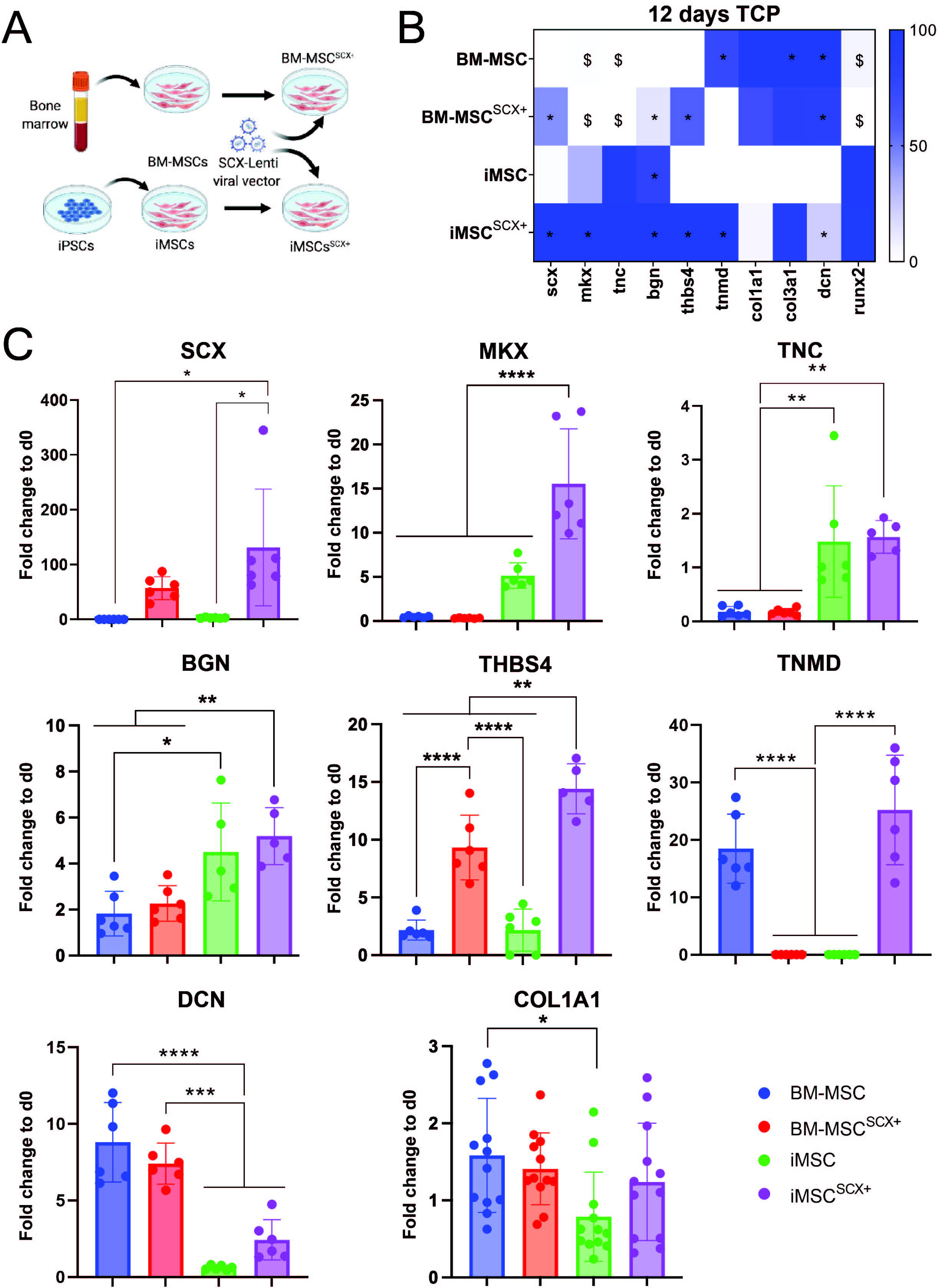
iMSC^Scx+^ show higher tenogenic potential than other cell types. **(A)** Graphical representation of the generation of four cell types that were used in the static culture differentiation experiments. BM-MSCs and iMSCs were transduced with lentiviral vectors to overexpress Scx stably. Cells (BM-MSCs, BM-MSC^SCX+^, iMSCs, and iMSC^SCX+^) were maintained in vitro for 12 days and then gene expression analyses were performed. **(B)** Tenogenic marker expression levels are displayed as heat maps; n=6, *p<0.05, (significant upregulation compared to baseline); ^$^p<0.05 (significant downregulation compared to baseline). **(C)** Gene expression levels of tenogenic markers were different between the four cell types after 12 days of culture in vitro. Between group comparisons were performed using one-way ANOVA with Tukey’s post-hoc; n=6, *p<0.05, **p<0.01, ***p<0.001, ****p<0.0001.

### 3.3 Static culture of BM-MSCs, BM-MSC^SCX+^, iMSCs, and iMSC^SCX+^ under differential substrate stiffness conditions

First, we examined the effect of differential substrate stiffness (E~ 2kPa, 32kPa versus standard TCP culture plates) on tenocyte marker expression (Col1a1, Col3a1, Tnmd, Thbs4, Mkx, and Alpl) of primary porcine tenocytes for a duration of 8 cell passages (Suppl. Fig. 2). Col1a1 gene expression was detected up to P8 in porcine tenocytes cultured on all types of surfaces, and there was a significant increase in Col1a1 in the early passages on all 3 substrates compared to baseline. Although Tnmd was slightly downregulated with passaging, it remained close to baseline in all three substrates up to P7 in which it dropped to 0 for E~2kPa and TCP, while it was still close to baseline for E~32kPa. However, at P8 its expression had dropped to 0 in all substrates. In the case of Thbs4 baseline expression was retained up to P5 for the two softer substrates and then dropped to 0, while on TCP it was retained up to P7. Alpl was significantly upregulated in TCP compared to both softer substrates and compared to baseline. Mkx was significantly downregulated after 1 passage in all substrates and remained at the same lower levels compared to baseline expression (at day 0) until the end of the study (data not shown). Lastly, Col3a1 expression was doubled compared to baseline in all substrates and remained constant until the end of the study. Next, the effect of substrate stiffness on the BM-MSC and BM-MSC^SCX+^ tenogenic differentiation potential was investigated. Tenogenic marker expression was increased in the TCP group. Scx was significantly higher in BM-MSC^SCX+^ cultured on TCP versus all other groups. Furthermore, Col1a1, Mkx, Thbs4, Bgn, and Dcn were significantly higher in BM-MSC^SCX+^ cultured on TCP compared to the two softer substrates (Supp. Fig. 3). Lastly, the effect of differential substrate stiffness on iMSC and iMSC^SCX+^ differentiation was assessed (Fig. 2A). Tnc and Bgn were significantly higher in iMSCs and iMSC^SCX+^ cultured on TCP compared to the lower stiffness substrates in iMSCs (Fig. 2B). Tenogenic markers Scx, Mkx, Tnmd, and Thbs4 were significantly upregulated in iMSC^SCX+^ cultured on TCP compared to iMSCs and iMSC^SCX+^ cultured in all other substrates. Lastly, Il-6, Tnfα, and Il-1β were significantly higher in iMSC^SCX+^ grown in 32kPa compared to all other groups, while Tnfα and Col3a1 were significantly higher in iMSC^SCX+^ grown in 32kPa and TCP compared to all other groups. Col1a1 was close to baseline expression levels in iMSC^SCX+^ cultured on TCP (data not shown). Zero net secretion of collagen was found in BM-MSCs, BM-MSC^SCX+^, iMSCs and iMSC^SCX+^ compared to baseline levels on the three different substrates after 12 days (data not shown).

**Fig. 2.**
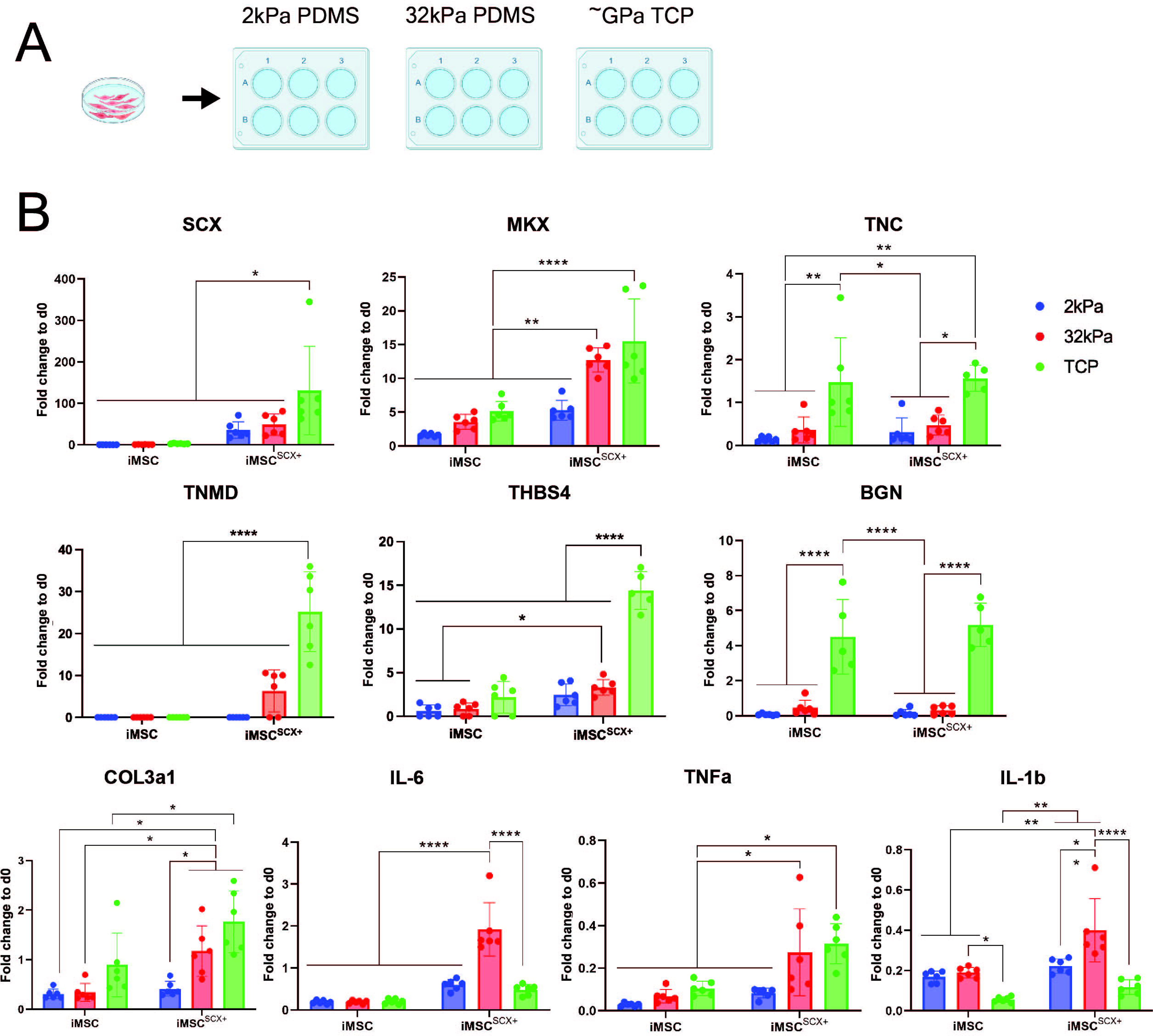
Lower substrate stiffnesses do not promote tenogenic differentiation in iMSCs with and without Scx. **(A)** Cells (iMSCs and iMSC^SCX+^) were differentiated in vitro for 12 days on surfaces with different substrate stiffnesses (2kPa, 32kPa, and TCP that is ~Gpa). (**B)** Gene expression analyses were performed. n=6 replicates/group. *p<0.05, **p<0.01, ***p<0.001, ****p<0.0001

### 3.4. Dynamic culture of iMSCs and iMSC^SCX+^ using cyclic uniaxial stretching

Tenogenic marker expression of iMSCs and iMSC^SCX***+***^ cultured on silicon plates in static conditions or cyclic stretching was analyzed at 3 and 7 days. Findings were summarized in a heat map showing changes compared to baseline (Fig. 3A). At 3 days, Scx, Mkx, and Bgn were upregulated in the iMSC stretched group compared to baseline and Scx and Mkx in the iMSC static group. At day 7, Scx, Col3a1, Pdgfra, Tppp3, Tnc, and Tnmd were significantly upregulated in the iMSC static group.

**Fig. 3.**
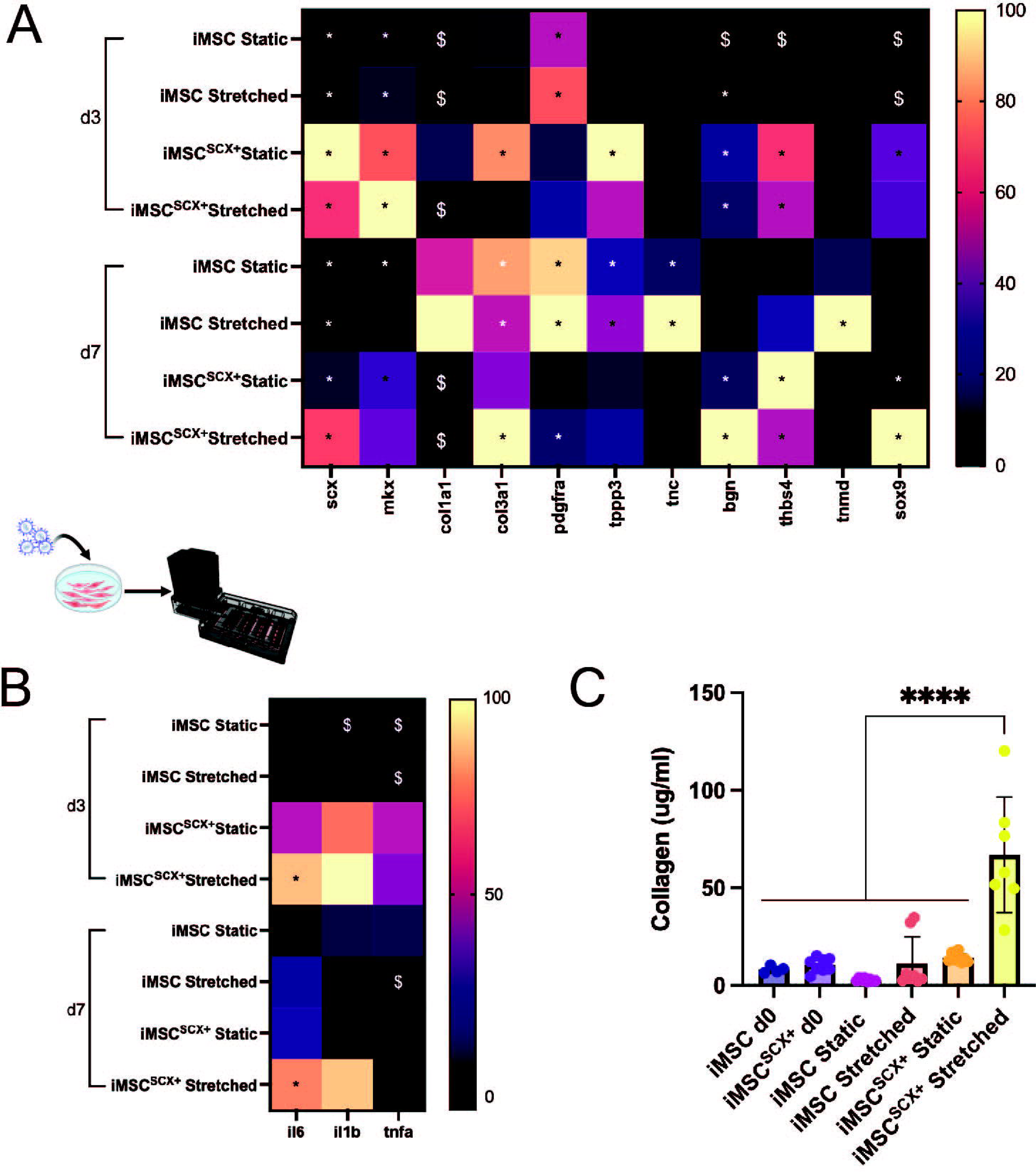
Tenogenic markers gene expression and matrix secretion are stimulated by Scx overexpression and uniaxial stretch in 2D bioreactor. iMSCs and iMSC^SCX+^ were stretched in a 2D bioreactor for 7 days. For static controls, cells were plated in matching plates with no stretching. (A) Gene expression analyses for iMSCs and iMSC^SCX+^ stretched for 3 days and 7 days in the 2D bioreactor. Tenogenic marker expression levels are displayed as heat maps; n=8, *p<0.05, (significant upregulation compared to baseline); ^$^p<0.05 (significant downregulation compared to baseline). (B) Inflammatory marker expression levels are displayed as heat maps; n=8, *p<0.05, (significant upregulation compared to baseline); ^$^p<0.05 (significant downregulation compared to baseline). (C) Collagen deposition following 7 days of stretch in the 2D bioreactor. n=8/group; *p<0.05, **p<0.01, ***p<0.001, ****p<0.0001.

Cyclic stretching of iMSC^SCX+^ for 3 days resulted in an upregulation of Scx, Mkx, Bgn, and Thbs4. At day 7, Thbs4 was still upregulated, Sox9 was also significantly upregulated, while Mkx dropped to baseline levels (Fig. 3A). Col3a1 was significantly upregulated after 7 days of stretching. Interestingly, in iMSC^SCX+^ baseline expression of Col1a1 and Col3a1 was significantly downregulated compared to iMSC baseline expression (Suppl. Fig. 4). Tnfα was significantly downregulated in iMSCs at both timepoints, while Il-6 was upregulated in iMSC^SCX+^ at both timepoints compared to baseline (Fig. 3B). Newly synthesized secreted collagen was significantly higher after 7 days of cyclic stretching in iMSC^SCX+^ compared to iMSCs that were also stretched for the same time, as well as both cell types that were cultured for 7 days in the silicone plates (static condition, Fig. 3C).

Cellular morphology and orientation of iMSCs and iMSC^SCX*+*^ was examined after 7 days of uniaxial loading or static culture via phalloidin staining of the actin fibers of the cytoskeleton. Actin filaments aligned perpendicular to the axis of the load in the stretched plates, whereas a stochastic cytoskeleton alignment was observed in the static culture (Fig. 4A). Additionally, an unbiased morphometric analysis was conducted on both culture conditions in iMSCs and iMSC^SCX*+*^ by using high content image analysis of the nuclei, whereby the operator of the software was blinded to the groups. Nucleic orientation assessment showed that uniaxial cyclic loading for 7 days significantly affected the frequency distribution of nuclear orientation in iMSCs and iMSC^SCX*+*^, which was assessed as an angle from the axis of loading (Fig. 4B). The medians of nuclear orientation of the static and the stretched conditions were shown to be significantly different. In iMSCs after 7 days of cyclic stretching, 54% of stretched nuclei were found at 65-90° angle compared to 33.7% nuclei in the static condition in the same range.

**Fig. 4.**
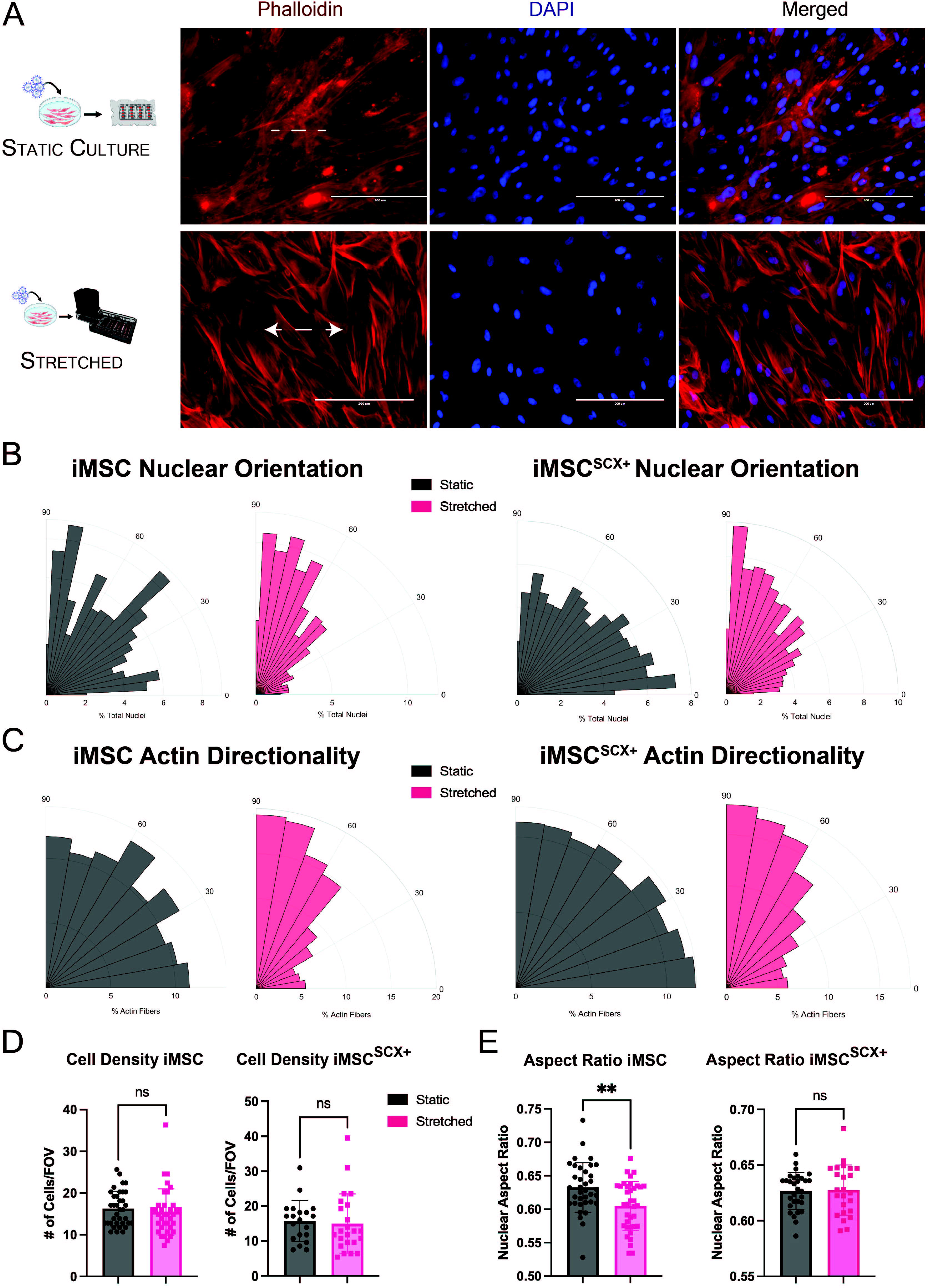
Cyclic stretching altered nuclear orientation and fiber alignment in iMSCs and iMSC^SCX+^. iMSCs and iMSC^SCX+^ were seeded on deformable silicone plates, stretched in a 2D bioreactor for 7 days (“stretched” condition), fixed with phalloidin for actin filaments (red) and counterstained with DAPI to mark nuclei (blue). Cells were also plated on similar plates that were placed in the incubator for the same period (static culture controls, “static”). **(A)** iMSC^SCX+^ are shown. White arrows display the direction of applied load. Scale bars are set at 200µm. **(B)** Unbiased morphometric analyses based on DAPI nucleic staining were performed to assess the effect of uniaxial cyclic loading on cellular orientation (n=7 individual wells/ group). The frequency distribution of nuclear orientation angles was plotted for the two conditions as rose plots (iMSCs left panel, iMSC^SCX+^ right panel). In iMSC^SCX+^ the distributions and their individual medians in the two tested conditions were significantly different and the stretched cells displayed skewness towards the 90°. **(C)** The frequency distribution of actin filament angles was plotted for iMSCs and iMSC^SCX+^. In iMSC^SCX+^ the two distributions were significantly different from each other and >50% of filaments of the stretched condition were found to be in 60-90°, whereas the static culture had a stochastic orientation of ~10% fibers found in each bin. n =7 wells/group **(D)** Cell density was computed by dividing the total number of nuclei by the area of each field of view. n=7/wells. **(E)** Nuclear Aspect Ratio (width to height of the nucleus) was computed to assess whether nuclei were elongated in each state. Values close to 1 reflect rounded nuclei, whereas smaller fractions represented more elongated nuclei, n=7 wells, **p<0.01.

Additionally, 36% of static culture nuclei were found 0-35°, unlike the stretched condition with 17.6% nuclei in the same range. (Fig. 4B; left panel). In iMSC^SCX*+*^ after 7 days, 42.5% of stretched nuclei displayed an angle of 65-90°, as opposed to only 26.5% of the nuclei of the static culture (Fig. 4B; right panel). In contrast, 30.2% of the static culture nuclei displayed an angle of 0-20°, while only 15.1% in the stretched plate were found in the same angular range (Fig. 4D). Furthermore, the frequency distribution of nuclear angles was skewed towards 90°, as opposed to the static culture, which showed a stochastic alignment of nuclei in all directions. Actin filaments displayed a different orientation distribution in the static culture versus the stretched condition in both iMSCs and iMSC^SCX^ after 7 days of cyclic stretching. In iMSCs that were stretched, 54.1% of actin fibers were found to be aligned in 60-90° angles compared to the axis of stretch, compared to 33.5% of the static condition. Similarly, in iMSC^SCX*+*^ in the stretched condition, >50% (50.8%) of fibers were found to be aligned in 60-90°, while in the static condition, there were similar fractions of fibers aligned in each directionality cluster. For both cell types, even though the two medians were not statistically different, the frequency distributions between the two conditions were shown to be significantly different. Cell density was similar for both conditions and cell types (Fig. 4D). Stretched iMSCs resulted in significantly lower aspect ratios (Fig. 4E). The frequency distribution of the nuclei in iMSC^SCX*+*^ showed that in both conditions, 76% of all nuclei had aspect ratios between 0.55-0.8 (data not shown) but the distributions or medians were not significantly different (Fig. 4E).

Further qualitative phalloidin staining of actin fibers and ICC against Tnmd and DAPI in iMSCs and iMSC^SCX+^ was performed (Fig.5). Following 7d of uniaxial stretching, quantitative assessment of median Tnmd intensity showed no difference between any of the groups for both cell types (data not shown). However, Tnmd staining appeared to be brighter and concentrated around nuclei in the stretched condition. In contrast, the static culture Tnmd staining displayed a more diffuse pattern for iMSCs and iMSC^SCX+^.

**Fig. 5.**
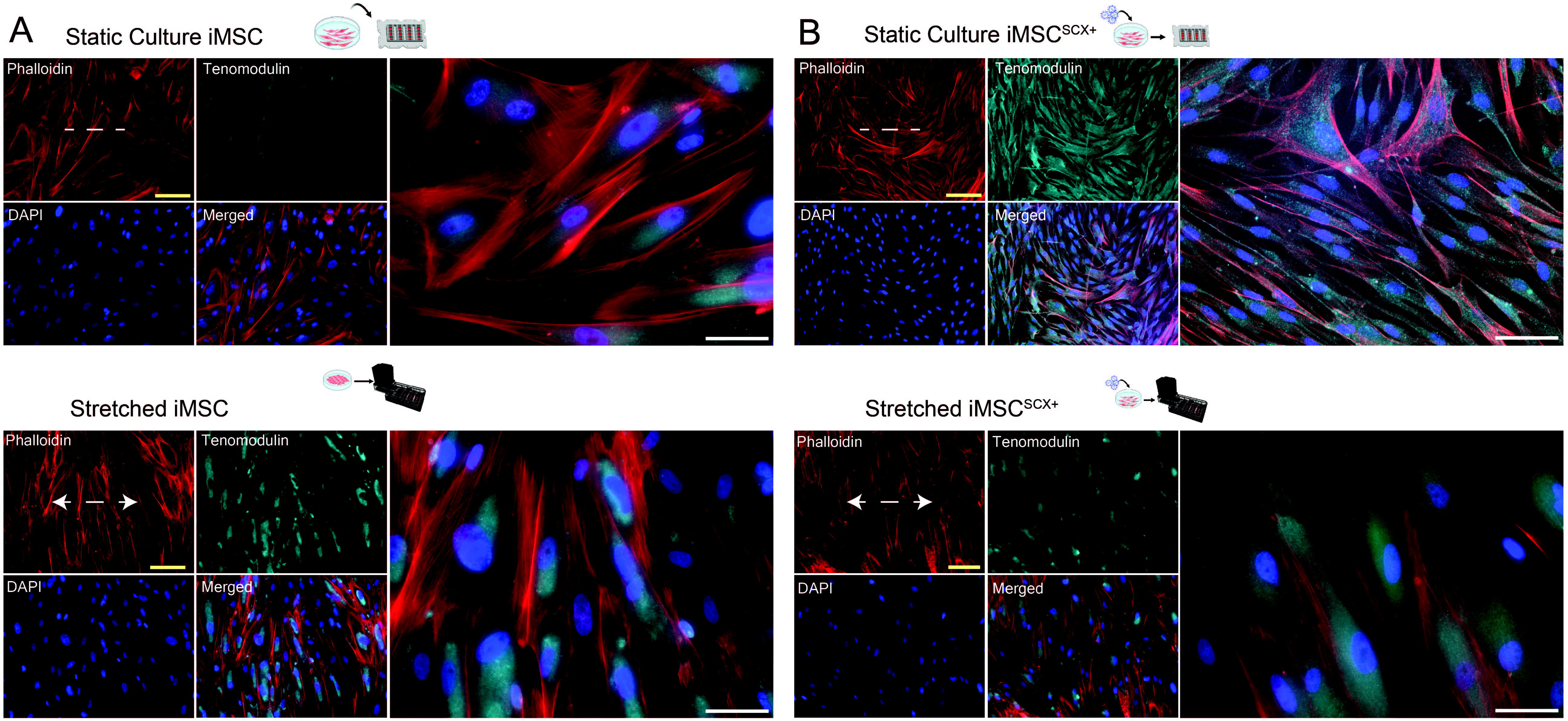
Confirmation of tenogenic differentiation on 2D bioreactor using immunocytochemistry. Cytoskeleton visualization using phalloidin staining of actin filaments (red), nucleic staining with DAPI (blue), and immunocytochemical staining for Tnmd (cyan) was performed in the static culture (“static”) and cyclic stretch (“stretched”) conditions in iMSCs **(A)** and iMSCSCX+ **(B).** Cells were seeded into silicone deformable plates with the same density. In the stretched condition, they were stretched uniaxially for 7 days in a 2D bioreactor, while for the static condition they were placed in the incubator for the same timeframe. Channels are shown individually and merged for the same field of view. A merged higher magnification image is also shown on the right for each cell type and condition. Dotted line displays the long axis of the silicone plate which was the axis of the longitudinal stretch that was applied with this bioreactor system. Two different magnifications are shown for the two conditions. Scale bars represent 130μm (yellow) and 60μm (white).

Finally, Collagen-1 staining was performed for iMSCs and iMSC^SCX+^ following 7d of cyclic loading and compared to the static condition (Fig. 6). For both cell types, Collagen-1 fiber deposition seemed to be in parallel to the actin fibers, unlike the static conditions, where the matrix displayed a more random pattern.

**Fig. 6.**
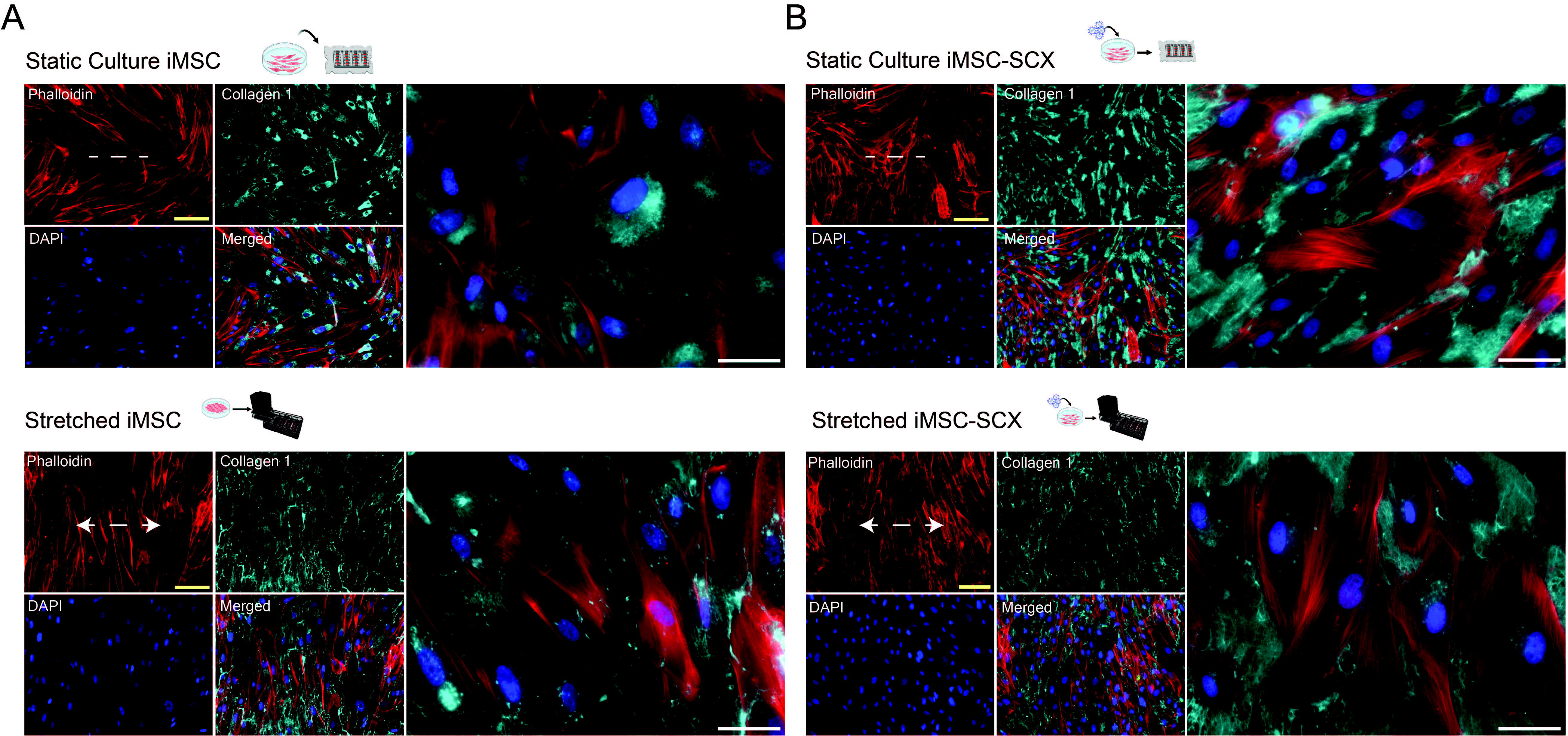
Immunocytochemistry for Collagen-1 and actin (phalloidin) following cyclic stretching in 2D bioreactor. Cytoskeleton visualization using phalloidin staining of actin filaments (red), nucleic staining with DAPI (blue) and immunocytochemical staining for Collagen-1 (cyan) was performed in the static culture (“static”) and cyclic stretch (“stretched”) conditions. Cells iMSCs **(A)** and iMSCSCX+ **(B)** were seeded into silicone deformable plates with the same density. In the stretched condition, they were stretched uniaxially for 7 days in a 2D bioreactor, while in the static they were placed in the incubator for the same timeframe. Channels are shown individually and merged for the same field of view. A merged higher magnification image is also shown on the right for each cell type and condition. Dotted line displays the long axis of the silicone plate which was the axis of the longitudinal stretch that was applied with this bioreactor system. Two different magnifications are shown for the two conditions. Scale bars represent 130μm (yellow) and 60μm (white).

## 4. Discussion

In the present study we investigated the effect of combined stable Scx overexpression and biomechanical stimulation to induce tenogenic differentiation of BM-MSCs and iMSCs. We demonstrated the following: (a) lentiviral vector-mediated stable overexpression induced tenogenic differentiation *in vitro* (Fig. 1); (b) when altering substrate stiffness, cell culture on softer substrates did not promote tenogenic differentiation of BM-MSCs or iMSCs compared to standard tissue culture polystyrene with and without Scx overexpression (Fig. 2); (c) uniaxial cyclic stretching was able to induce upregulation of tenogenic markers in both iMSCs and iMSC^SCX+^ at day 7, with significantly higher levels of expression in the iMSC^SCX+^ group (Fig. 3); (d) uniaxial stretching resulted in changes in nuclear orientation and cytoskeleton alignment in iMSCs and iMSC^SCX+^ cells (Fig. 4); and (e) Scx overexpression in iMSCs combined with stretching resulted in Collagen-1 fiber alignment and Tnmd deposition, indicating complementing effects of biological and mechanical cues to induce tenogenic differentiation in iMSC^SCX+^ compared to iMSCs (Fig. 5–6). To our knowledge, this is the first study to demonstrate tenogenic differentiation processes in iPSC-derived MSC^SCX+^ under 2D cyclic tension. Scx stable overexpression resulted in a significant upregulation of earlier and later tendon markers *in vitro*, especially in iMSC^SCX+^. These findings are in line with prior literature reporting that Scx is an important transcription factor that regulates syndetome specification and tendon formation.^34; 47^ Specifically, we demonstrated that BM-MSC^SCX+^ cells significantly upregulated Bgn, Thbs4, and Dcn after 12 days of culture compared to their own baseline. Compared to naïve BM-MSCs, Thbs4 was increased and Tnmd decreased in BM-MSC^SCX+^ cells (Fig. 1C). In contrast to our study, Alberton *et al.* demonstrated Scx transduction of the SCP-1 hTERT immortalized BM-MSC cell line to induce Tnmd expression in a static *in vitro* culture system.^21^ These differences may be a result of the cell source, since they used an immortalized cell line while our study tested primary low passage BM-MSCs.^21^

In the current study, iMSC^SCX+^ showed a significant upregulation in a larger number of tenogenic markers, including Tnmd as well as Mkx, Tnc, Bgn, and Thbs4 compared to baseline and the other cell types that were assessed. The stronger response of the iMSCs to Scx overexpression might be due to differences in the cell origin: while BM-MSCs are obtained from different donors of different ages displaying phenotypic variability that are known to potentially affect their *in vitro* performance, iMSCs are considered a more homogeneous population of cells, since they are generated and expanded from a single origin.^48^ Moreover, iMSCs are derived from embryonic-like iPSCs and were reported to have a rejuvenation gene signature.^49^ In addition, iMSCs were shown to have superior survival and engraftment after transplantation,^50^ higher proliferative capacity, lower senescence rate,^51^ and higher immunomodulation ability^51; 52^ compared to BM-MSCs.

The effect of substrate stiffness on the differentiation of multiple adult stem cells has been studied extensively.^53–55^ Mechanovariant substrates and selected ligands have been used in the past as a strategy to achieve directed differentiation of BM-MSCs towards tendon.^56; 57^ In the present study, use of mechanovariant substrates (stiffnesses of E~2kPa, 32kPa and ~GPa) did not promote expression of tenogenic differentiation markers compared to culture on standard TCP in BM-MSC and iMSC with and without Scx overexpression after 12 days of static culture (Fig. 2B). These results are in contrast with a previous study using BM-MSCs cultured at E ~40kPa for 2 weeks, showing an upregulation of Scx, Tnmd, and Tnc.^56^ Differences between the results may be due to differences in donor or method of cell expansion prior to the experiments.^58^ For example, Li *et al* found that priming of primary rat BM-MSCs on a stiff surface resulted in mechanical memory that lasted for more than 2 weeks through a micro-RNA-21-controled mechanism.^59^ They showed that miR-29 expression was established after mechanical priming for 3 passages and this effect could be maintained for at least two passages after switching to a substrate with a higher or lower stiffness.^59^ The cells that were used for our experiments, BM-MSCs and iMSCs, were initially expanded on TCP plates (up to P4). Therefore, it is likely our initial expansion cycles on TCP could have primed the cells towards a stiffer phenotype that masked the effect of the softer substrates over the course of 12 days of our *in vitro* static culture. Since expansion and culture conditions differ widely between laboratories, the additional factor of surface stiffness used to expand cells may prove to be an important consideration in experimental designs and in the methodologies used in tendon cell therapy applications and it should be addressed in future studies.

Tenocytes and tendon stem/progenitor cells dedifferentiate when cultured statically in regular tissue culture plates.^7^ Tnmd was immediately downregulated with *in vitro* culture of pig tenocytes (Supp Fig. 2), as has been reported.^7^ Interestingly, Alpl expression was close to baseline for the softer substrates, but significantly upregulated in TCP, which may indicate osteogenic differentiation of cultured tenocytes over time on more rigid substrates.^60^ This suggests that expansion of tenocytes and iPSC-derived tenocytes for cell therapy applications may need to be performed in softer substrates in order to maintain their phenotype prior to use.

In the present study, we stimulated iMSCs with 4% uniaxial strain and 0.5Hz for 2 h/day in a 2D bioreactor using deformable silicone plates, which is in line with other studies showing that physiologically relevant loads range from 4%-6% strain at the cellular level of tendon.^29^ The final seeding density was chosen to avoid monolayer overgrowth on the silicone plate, which has been described to contribute to static tension.^61^

After 7 days of cyclic mechanical stimulation, a significant upregulation of Scx, Tnmd, Tnc, Col3a1, Tppp3 as well as Pdgfra in stretched iMSCs was detected (Fig. 3A). Using the same bioreactor system (CellScale MCFX), but applying higher total strain amplitude and duration (10% at 1Hz, for 12h/day), Gaspar *et al.* were not able to detect changes in gene expression of Scx, Tnmd, and Col1a1 and collagen deposition in BM-MSCs and human tenocytes.^61^ However, they reported significant upregulation of Thbs4 after 3 days. This could reflect the higher strains that were applied in that study, since BM-MSCs aligned parallel to the load which is in contrast with a big body of research showing perpendicular cell alignment with physiological strains.^62^ In our study, stretched iMSC^SCX+^ significantly upregulated Scx, Col3a1, Bgn, Thbs4, Sox9, and Pdgfra, but not Tnmd at day 7 (Fig. 3A). Comparable to our study, Chen *et al*. demonstrated an induction of Col1a1, Col1a2, Col14, and Tnmd and increased ECM deposition in lentiviral-mediated Scx-overexpressing hESC-MSCs that were assembled in multilayered cell sheets and cultured under uniaxial dynamic cyclic load (10% strain, 1Hz, 2 h/day) for up to 21 days.^63^ Implantation of these cells resulted in an improved tendon regeneration in a small-size tendon defect in a nude mouse model.^63^

Assessment of new ECM deposition displayed significantly higher total collagen content in the iMSC^SCX+^ cells stretched for 7 days compared to iMSCs in both static and stretched conditions and to iMSC^SCX^ in the static culture at the same timepoint (Fig. 3C). This indicates a potential synergistic effect of Scx overexpression and uniaxial loading resulting in not only tendon marker expression, but also more importantly secretion of ECM proteins to the media. BM-MSC^SCX+^ in static culture and cell sheets of Scx^+^ human ESCs-derived cells under loading have been shown to result in increased collagen deposition.^21; 63^ Lastly, Il-6 was significantly upregulated after 12 days of static culture in iMSC^SCX+^ cultured on the 32kPa substrate but not in any other groups (Fig. 2). Additionally, Il-6 was upregulated after 3d and 7d of uniaxial stretching in iMSC^SCX+^ but not in iMSCs (Fig. 3B). Inflammatory markers including Il-6 have been shown to be upregulated by loading of higher magnitude (8-12%) in 2D uniaxial stretching systems^29^ and its upregulation has been reported in tendinopathies and ruptured tendons.^64^ However, in this work, Il-1β and Tnfα did not change compared to baseline in iMSCs and iMSC^SCX+^ cultured on the three substrates, nor when they were stretched in the 2D bioreactor (Fig. 3B). Il-6 was also not changed following stretching in iMSCs (Fig. 3B). In human tendons, Il-6 is upregulated following exercise, and it is a potent stimulator of collagen synthesis.^65^ Thus, Il-6 upregulation might have contributed to the increased ECM deposition in stretched iMSC^SCX+^ (Fig. 3B-C).

The distribution of nuclear and cytoskeleton orientation was very distinct between the two conditions, suggesting that the cells re-oriented as a response to the stretch stimulus (Fig. 4B-C). While the statically cultured cells displayed stochastic nuclear arrangement in all directions, in the stretched cells the predominant orientation of nuclei was perpendicular to the applied load. Consistent with our findings, previous reports have shown cell orientation perpendicular to the stretching direction when physiological strains (≤~7%) were applied.^62; 66^ Nucleic aspect ratios (width to height) of stretched versus static cells were unaltered in iMSC^SCX+^ but significantly lower in stretched iMSCs compared to the static condition, pointing to more elongated nuclei (Fig. 4E).

Actin filament staining and ICC for Tnmd in iMSCs and iMSC^SCX+^ static and stretched cells showed presence of Tnmd in the entire cytoplasm (Fig. 5). Tnmd seemed more concentrated around the nuclei in the stretched group for both cell types. Intriguingly, Tnmd gene expression was unchanged at days 3 and 7 with cyclic stretching in iMSC^SCX+^ cells (Fig. 3 and Suppl. Fig.4). Tnmd is directly regulated by Scx,^67^ and previous studies using Scx overexpression in BM-MSCs have reported Tnmd upregulation.^21^ However, recent findings using equine ESCs and fetal tenocytes to overexpress Scx have shown Tnmd gene downregulation, suggesting that mRNA and protein levels are differentially regulated and therefore need to be assessed in tandem.^68^ Lastly, Collagen-1 staining in iMSCs and iMSC^SCX+^ that were loaded for 7 days (Fig. 6) showed a more organized collagen fiber network which was in parallel to the actin fibers and perpendicular to the axis of stretch, unlike the static groups, which displayed a more random orientation of fibers. Complemented with our collagen assay data, it can be concluded that iMSC^SCX+^ that were cyclically loaded for 7 days, resulted in a newly formed ECM fiber network whose orientation might have been instructed by mechanical regulation.

This study is not without limitations. It could be postulated that even though our transduction efficiency was close to 100%, there may have been cells that were not transduced with Scx. Thus, the proportion of Scx^+^ cells may decrease with expansion and some of the transduced cells might have lost Scx expression due to gene silencing.^69^ Cell sorting could potentially be used to obtain a purer iMSC^SCX+^ population. Although, cell sorting was not necessary in this study, since we were able to achieve iMSC^SCX+^ tenogenic differentiation, it may still be needed for future *in vivo* applications. Further, it has been postulated that the nominal strain applied in a 2D stretching system may be higher compared to the actual strain experienced by the cells.^70^ The latter may vary depending on bioreactor system, cell density and cell type (size, morphology, etc.), which hampers comparison and standardization between different studies.^29; 61^ To this end, we performed a series of pilot studies to optimize the stretch protocol and used complementary morphometric analyses to ensure we were stretching the cells within their physiological range. Lastly, the lack of adequate specific markers to assess temporal tenocyte gene expression changes has resulted in difficulty to characterize differentiation efforts and to discriminate between the effects of biological cues versus biomechanical ones that are crucial for tendon tissues.^25^ Therefore, we need additional functional assessments to understand changes following differentiation efforts and the effects of mechanical stimulation on the maturation of the cellular phenotype.

## Conclusions

An appropriate and potent cell therapeutic candidate for tendon regeneration is needed to address today’s challenges in tendon and ligament repair. Our data provide evidence that iMSCs, genetically engineered to express Scx and stimulated by cyclic mechanical stretch to drive the cells into iTenocytes, may be a potential candidate for tendon cell therapy applications. Tenogenic potential of iMSC^SCX+^ was demonstrated by upregulation of early and late tendon markers, increased collagen deposition and ECM alignment, Tnmd expression, morphometric and cytoskeleton-related changes. Overall, iMSCs performed better than BM-MSCs, which might be due to the fact that iMSCs are developmentally more plastic.

In future research, the functional impact of iTenocytes on tendon and ligament repair should be evaluated in an animal tendon injury model, providing an environment of physiological locomotion.

## Acknowledgments

This study was supported by the NIH/NIAMS K01AR071512 to DS. The two lentivirus packaging plasmids were a gift by the Simon Knott laboratory (Department of Biomedical Sciences, Cedars-Sinai). The authors would like to thank Ms. Catherine Bresee (Cedars-Sinai Biostatistics Core Manager) for consultation on statistical analyses performed to compare the frequency distributions generated in the morphometric analyses.

## Disclosure of Potential Conflicts of Interest

The authors have no conflict of interest.

## Notes

### Competing Interest Statement

The authors have declared no competing interest.

