## Supplement Tables 1-2, Figures 1-4 for "Directing iPSC Differentiation into iTenocytes using Combined Scleraxis Overexpression and Cyclic Loading"

**Suppl. Table 1**

A. **Human Primers used for gene expression analysis**

| **Gene** | **Full Name** | **Pubmed RefSeq** | **Assay ID/Cat#** |
| --- | --- | --- | --- |
| Scx | Scleraxis | NM_001080514.2 | Hs03054634_g1 |
| Mkx | Mohawk | NM_001242702.1 | Hs00543190_m1 |
| Col1a1 | Collagen type I alpha 1 chain | NM_000088.3 | Hs00164004_m1 |
| Col3a1 | Collagen type III alpha 1 chain | NM_000090.3 | Hs00943809_m1 |
| Egr1 | Early growth response 1 |  | Hs00152928_m1 |
| Tnmd | Tenomodulin | NM_022144.2 | Hs00223332_m1 |
| Tnc | Tenascin C | NM_002160.3 | Hs01115665_m1 |
| Dcn | Decorin | NM_001920.4 | Hs00370384_m1 |
| Thbs4 | Thrombospondin 4 | [NM_001306212.1](http://www.ncbi.nlm.nih.gov/nuccore/NM_001306212.1) | Hs00170261_m1 |
| Bgn | Biglycan | NM_001711.5 | Hs00959143_m1 |
| Alpl | Alkaline phosphatase, biomineralization associated | NM_000478.5 | Hs00758162_m1 |
| Runx2 | Runt-related transcription factor 2 | NM_001015051.3 | [Hs01047975_m1](https://www.thermofisher.com/taqman-gene-expression/product/Hs01047975_m1?CID=&ICID=&subtype=) |
| Spp1 | Secreted phosphoprotein 1 | NM_000582.2 | Hs00959010_m1 |
| Sparc | Secreted protein acidic and cysteine rich | NM_001309443.1 | Hs00234160_m1 |
| Itga5 | Integrin subunit alpha 5 | NM_002205.4 | [Hs01547673_m1](https://www.thermofisher.com/taqman-gene-expression/product/Hs01547673_m1?CID=&ICID=&subtype=) |
| Itgb2 | Integrin subunit alpha 2 | NM_000211.4 | [Hs00164957_m1](https://www.thermofisher.com/taqman-gene-expression/product/Hs00164957_m1?CID=&ICID=&subtype=) |
| Il-6 | Interleukin 6 | NM_000600.4 | HS00985639_m1 |
| Il-1b | Interleukin 1 beta | NM_000576.2 | HS01555410_m1 |
| Tnfa | Tumor necrosis factor alpha | NM_000594.3 | Hs00174128_m1 |

**B. Pig Primers used for gene expression analysis**

| **Gene** | **Full Name** | **Pubmed RefSeq** | **Assay ID/Cat#** |
| --- | --- | --- | --- |
| Thbs4 | Thrombospondin 4 | XM_021084490.1 | [Ss06910153_m1](https://www.thermofisher.com/taqman-gene-expression/product/Ss06910153_m1?CID=&ICID=&subtype=) |
| Tnmd | Tenomodulin | NM_001099934.1 | Ss03389154_m1 |
| Col1a1 | Collagen type I alpha 1 chain | XM_021067153.1 | Ss03373340_m1 |
| Col3a1 | Collagen type III alpha 1 chain | NM_001243297.1 | Ss04323794_m1 |
| Alpl | Alkaline phosphatase, biomineralization associated | XM_021097681.1 | Ss06879568_m1 |
| Bgn | Biglycan | XM_003135475.5 | Ss03375454_u1 |
| Mkx | Mohawk | XM_021064711.1 | Ss06835432_m1 |

**SUPPLEMENTAL FIGURES**

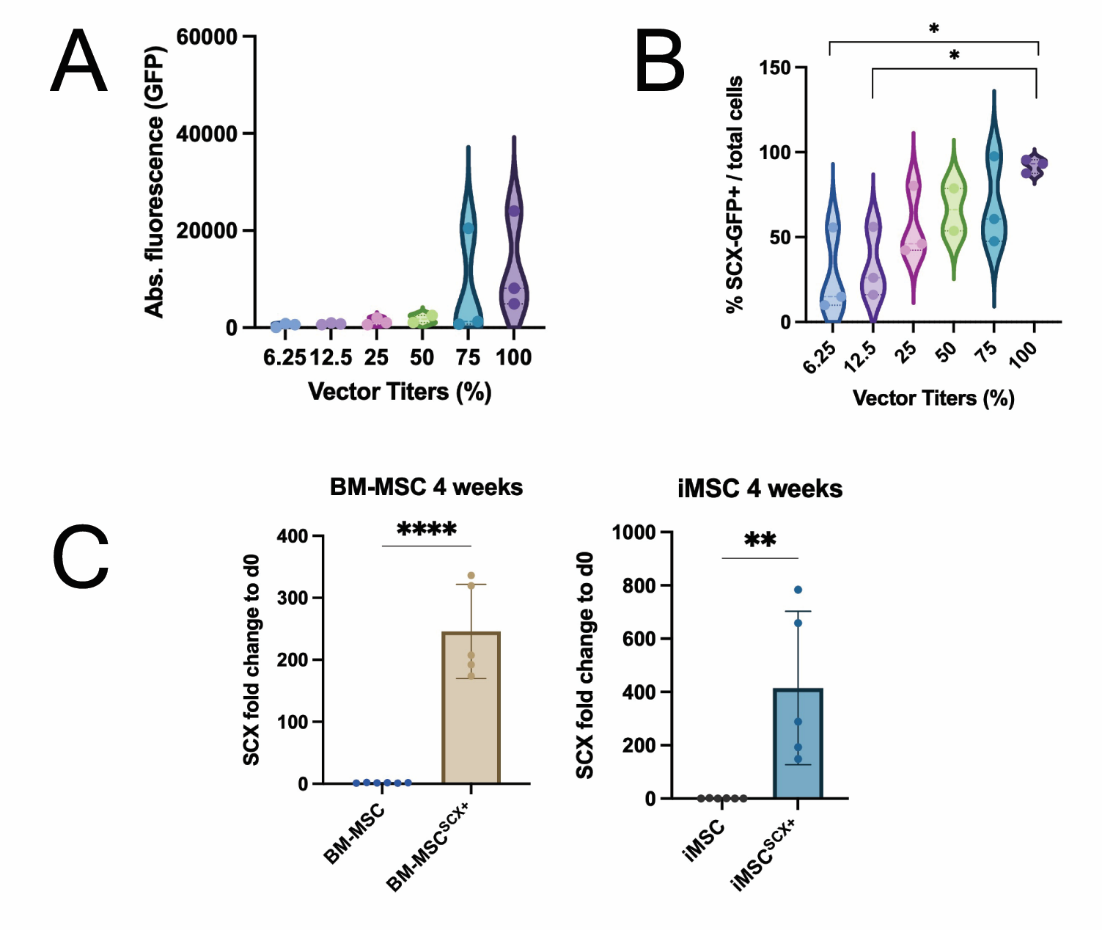

**Suppl. Fig. 1. Flow cytometry was used to assess transduction efficiency with a 2^nd^ generation lentivirus vector.** *Vector titers were validated by flow cytometry analysis for all cells. Representative transduction efficiency analyses conducted on BM-MSCs.* ***A).*** *Absolute fluorescence per virus titer.* ***B).*** *Transduction efficiency evaluated as % percentage of SCX-GFP^+^ cells. n=3 independent transductions.* ***C).*** *Gene expression analysis performed on BM-MSCs and BM-MSC^SCX+^ (left) and iMSCs and iMSC^SCX+^(right) that were cultured for 4 weeks. Scx was significantly upregulated in cells that were transduced with the Scx vectors. p was set at<0.05 **** p <0.0001, ** p<0.01*

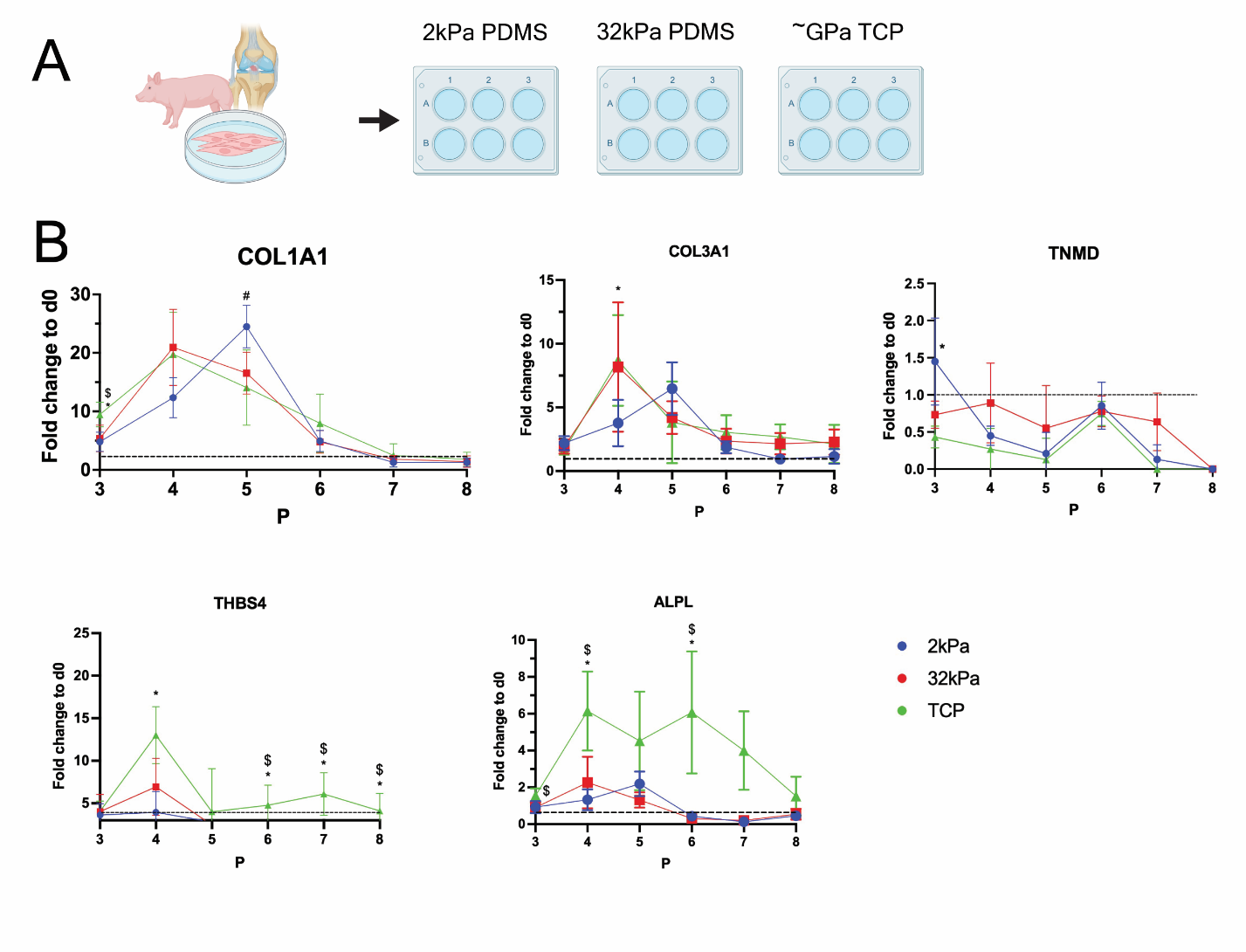

**Suppl. Fig. 2.** **Assessment of the effect of differential substrate stiffness on the retention of pig tenocyte phenotype over time**. ***A).*** *Pig tenocytes were cultured until P8 in three different substrates (2kPa, 32kPa, and TCP).* ***B).*** *Gene expression analysis was performed for Col1a1, Col3a1, Tnmd, Thbs4, and Alpl. A mixed repeated measures model was used to compare the three groups, with Tukey’s post hoc corrections for between group comparisons at each passage. *p<0.05 2kPa vs. TCP, $ p<0.5 32kPa vs. TCP, # p<0.05 2kPa vs. 32kPa*

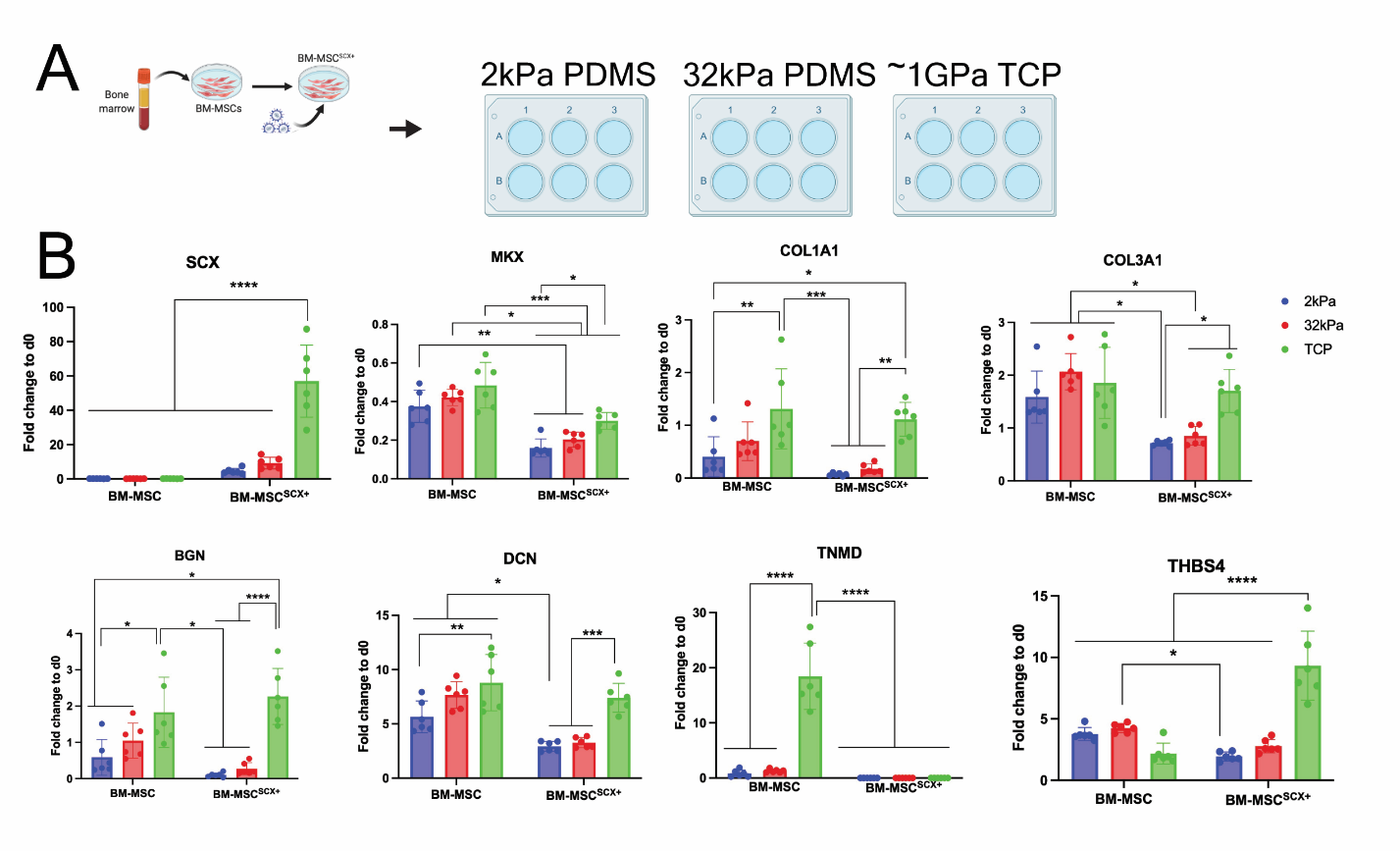

**Suppl. Fig. 3. Softer surface does not promote tenogenic differentiation in BM-MSCs with and without Scx,** ***A).*** *Cells (BM-MSCs and BM-MSC^SCX+^) were differentiated in vitro for 12 days on surfaces with different cell substrate stiffnesses (2kPa, 32kPa, and TCP that is ~GPa) and then* ***B).*** *gene expression analyses were performed.* *p<0.05, **p<0.01, ***p<0.001, ****p<0.0001

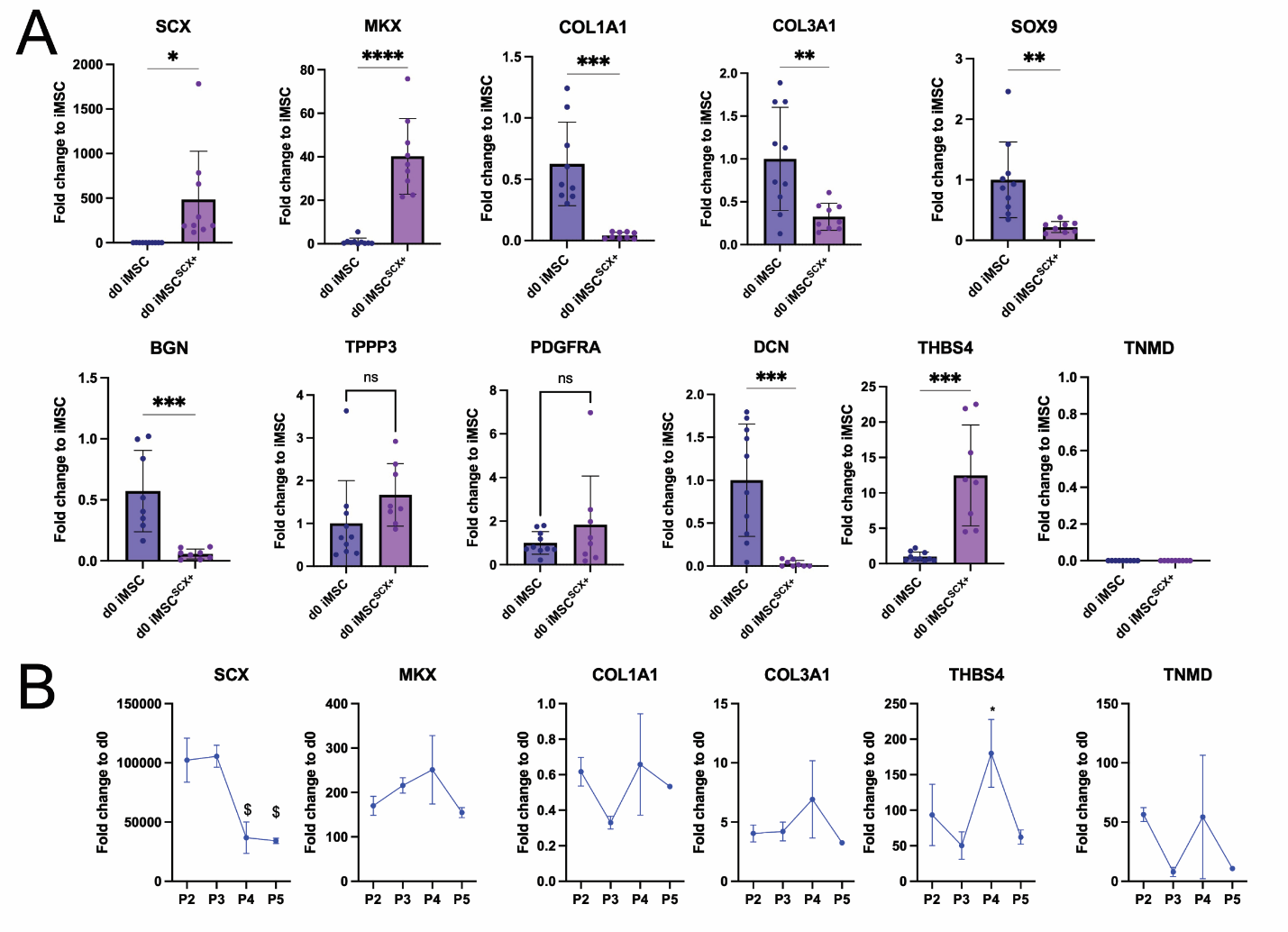

**Suppl. Fig. 4.** **Gene expression of tenogenic markers in iMSCs and iMSC^SCX+^***.* ***A).*** *Baseline expression between iMSCs and iMSC^SCX+^ that were used in the experiments of this work. Several relevant markers are displayed.* ***B).*** *Gene expression of tenogenic markers over time in iMSC^SCX+^. *p<0.05, (significant upregulation compared to baseline); ^$^p<0.05 (significant downregulation compared to baseline).*
